# Categorical rhythmic patterns do not predict song preference in zebra finches

**DOI:** 10.64898/2026.10.05.756641

**Authors:** Marco Maiolini, Michelle Spierings

## Abstract

Categorical rhythmic patterns are fundamental features found in language and music across cultures. In humans, these rhythmic patterns occur in both vocal production and perceptual preference, highlighting a link between rhythmic production and preference. Categorical rhythmic patterns have recently been identified in animal vocalisations too, but whether rhythm production is linked with rhythmic preferences in non-human animals remains unclear.

Within this context, zebra finches (*Taeniopygia castanotis*) are of particular interest, as both sexes develop a song preference, and their song has been suggested to exhibit an isochronous temporal structure. Here, we tested for the first time whether zebra finches prefer songs containing discrete rhythmic patterns, using operant preference tests with either two or four rhythmic patterns.

Our results show that no categorical rhythmic pattern was consistently preferred across the population, although responses varied among individuals. These findings indicate that categorical rhythmic patterns are unlikely to be a primary driver of song preference in zebra finches. Together with previous work in other vocal-learning birds, our results suggest that preferences for categorical rhythmic patterns may not be universal and could instead reflect species-specific features of vocal displays rather than the rhythmic categories characteristic of human music.

## Introduction

Rhythm, defined as the way elements occur over time [1], is an important feature of interactive communication systems across both visual and acoustic modalities [2,3]. Categorical rhythmic patterns, which embody defined classes of intervals described by ratios of integers (e.g. 1:3, 4:7, 2:1), are one of the distinctive characteristics that shape music across cultures [4]. The presence of categorical rhythmic patterns in music across cultures gives rise to two different hypotheses. On the one hand, categorical rhythmic patterns could be the byproduct of the motor system [5, 6] and neural mechanisms [7–9], as they are relatively easy to perceive and produce [10]. On the other hand, rhythmic patterns may have emerged to facilitate the synchronisation and coordination between individuals, as evidenced by joint rushing activities [11,12] and work songs [13,14]. Overall, both hypotheses highlight the widespread importance of categorical rhythmic patterns in humans, which may have arisen from either social contexts or as mechanical byproducts.

Different categorical rhythmic patterns could be defined depending on which ratio of integers is embedded in the sequence. These patterns can be further classified as aperiodic, characterised by irregular distribution of events over time, or periodic, characterised by the regular recurrence of one or more temporal intervals. Among periodic rhythms, isochrony, equal temporal spacing between consecutive events, represents the simplest rhythmic structure. In contrast, patterns containing more than one interval duration are classified as heterochronous and are categorised into simple heterochronous rhythmic patterns with small integer ratios (e.g. 2:1:2:1; or 1:3:1:3…), and complex heterochronous rhythmic patterns, with complex integer ratios [3] (e.g. 7:4:7:4; or 13:9:13:9).

Among these different patterns, isochrony represents one of the core features of human rhythm production, being a shared component of music across cultures [4]. Together with simple heterochronous patterns, it emerges as a shared tendency in cross-cultural iterative studies, where participants repeatedly reproduce rhythms learned from others [10]. The relevance of rhythmic structure for humans goes beyond music production, extending to perception as well. For example, the processing of rhythmic patterns activates reward-related neural systems [15], and people tend to spend more time in an area with periodic rhythms rather than aperiodic rhythms [16].

Over the last decade, an increasing number of studies have examined the presence of rhythmic patterns in animal vocalisations [17]. Among the categorical rhythmic patterns found in humans, isochrony has been reported in the majority of species investigated, from insects [18], to fishes [19], anurans [20], birds [21,22] and mammals [23-30]. Furthermore, isochronous patterns have been found in all the developmental stages of indris’ song [31], suggesting isochrony as a conserved biological constraint. On the contrary, while heterochronous rhythmic patterns are commonly found in human music, they are rare in animal vocalisations, with simple heterochronous patterns being found only in the lemur Indris’ song [32,33]. Thus, although rhythmic structure is widespread in animal communication [34, 35], few studies have investigated whether rhythmic patterns in vocal production align with species-specific rhythmic preferences.

However, in order to form preferences for particular rhythmic patterns, animals need to be able to recognise and discriminate between different patterns. Studies on rhythm perception have addressed this question in primates, birds, and rodents. Among primates, rhesus macaques (*Macaca mulatta*) discriminate between isochronous and aperiodic rhythmic patterns [36, 37]. Similarly, rats (*Rattus norvegicus*) discriminate between periodic and aperiodic rhythmic patterns [38]. In birds, both budgerigars (*Melopsittacus undulatus*) [39] and zebra finches (*Taeniopygia castanotis*) [39,40] successfully discriminate between isochronous and aperiodic strings. These studies show only limited generalisation to the same rhythmic patterns with tempo or note alterations, suggesting that they might focus on local aspects of the sounds rather than the holistic pattern. Discriminating between rhythmic patterns reflects the ability to recognise differences between stimuli, which may evoke the development of a preference for one rhythm or a class of rhythms.

To date, rhythmic preference in animals has been investigated in only a few species. A study in mice by Perrodin and colleagues [41] examined which acoustic features make male song attractive. In the study, the female mice were placed in a two-compartment box and were exposed to different sound playbacks in each compartment. Females did not show preferences based on spectral manipulations, but spent more time in the compartment with original songs compared to temporally irregular songs. However, females did not show a preference when exposed to original songs and experimentally enhanced super-regular songs, suggesting avoidance of temporal irregularity but not necessarily a stronger preference for increased regularity. Importantly, because the rhythmic structure of the original songs was not described, it remains unclear whether females responded to periodicity itself or to specific embedded categorical rhythms.

A similar paradigm was used to test preferences for sounds with periodic versus aperiodic rhythmic beat patterns in humans and budgerigars [16]. The subjects were placed for 5 minutes in a preference test chamber with a speaker and a playback stimulus in each of the two sections. Humans showed a preference for periodic rhythmic stimuli, whereas budgerigars did not show a preference on the group level. However, female budgerigars showed a preference for the periodic rhythmic patterns, whilst the males showed an avoidance of them. As the aim of the study was to inspect rhythmic preferences fitting an underlying beat, the stimuli used were musical strings fitting periodic and aperiodic beat patterns, while the categorical rhythmic patterns in the stimuli were not described. Therefore, as in the mice study, it is challenging to interpret whether the preference shown was for the periodicity of the beat or the different embedded categorical rhythms nested in the musical string. Taken together, these studies do show that the temporal information in both natural vocalisations and artificial stimuli can influence auditory preferences.

One of the species that can discriminate between isochronous and aperiodic rhythmic patterns [39], but whose preference for it has never been investigated, is the zebra finch. Zebra finches are songbirds, closed-ended vocal learners with a complex fission-fusion society [42, 43]. In this species, only the males sing, with the juveniles learning the song from an adult male during a critical period [44]. In this process, the juvenile creates his unique song, sharing features with the tutor both in the spectral [45] and temporal domain [46]. The zebra finch song consists of a few introductory notes followed by the motif, which is repeated 2-3 times [47,48]. The motifs are formed by 4-7 syllables, which are constituted by smaller units, the elements [47,48]. Two studies have inspected the overall rhythm of the song, reporting a metronome-like pattern fitting with the isochronous rhythmic pattern [46,49-50]. Together, these features make zebra finches a strong model to investigate whether rhythmic preferences are aligned with categorical rhythmic patterns in production.

Here, we investigate whether zebra finches show species-specific preferences for categorical rhythmic patterns using an operant preference setup, where the birds voluntarily trigger playback sounds. Specifically, we test behavioural preferences for isochronous versus aperiodic rhythmic patterns, and we extend this preference test to include two different simple heterochronous rhythmic patterns.

## Methods

### Subjects

Thirty-one zebra finches participated in the two preference experiments (19 females and 12 males). Two birds (one female and one male), already familiar with the setup, were used as guide birds during the habituation phase. The guide birds landed on the operant perches, and naïve birds followed them to maintain proximity to conspecifics, thereby facilitating exploration of the perches and increasing perch landing. All the individuals were adults ranging from 1 to 6 years old at the time of the experiment. All the tested birds were naïve to the operant setup and the stimuli. When not tested, the birds were kept in single-sex group aviaries (2 x 2 x 1.2 m) at Leiden University under 21.5 °C and a 13.5L:10.5D cycle, with *ad libitum* access to food and water.

### Apparatus

The current study used an operant preference test to determine if zebra finches prefer particular rhythmic patterns. The setup consisted of an octagonal arena (85 cm diameter x 1 m high) with a perch positioned at 20 cm height on each side of the octagon (modification of setup from previous studies [51-53]). Food and water were provided *ad libitum* in the centre of the octagon arena at a height of 30 cm. Among the 8 perches present in the arena, which were equidistant and equally reachable from the centre, 4 constituted the operant perches. Whenever a bird landed on an operant perch, a sound playback from a nearby speaker (BLAUPUNKT CB 4500) was played at 70 dB SPL (Figure 1A). The sound was played once and continued until the end of the stimulus each time the perch was activated. All the occurrences were automatically recorded by a Python Software Program (4CTv1.0; 54). However, to avoid triggering an event due to hopping and to encourage the exploration of the setup, a delta time of inactivity of the perch was programmed at 250 milliseconds. An automatic software function enabled switching the sound associated with the perch at a specified time, allowing disentanglement of positional preferences from sound preferences. The operant setup stood in the middle of a sound-attenuated square room (3 x 3 x 2.4 m) at 75 cm height, illuminated by a fluorescent light (Philips MASTER TL5 HO) that emitted a daylight spectrum at a 13.5L:10.5D regime. The room was accessible through a door positioned in the corner between perches 1 and 2. An extra speaker underneath the operant setup (JBL Clip+) played colony sounds for the whole duration of the testing period.

**Figure 1.**
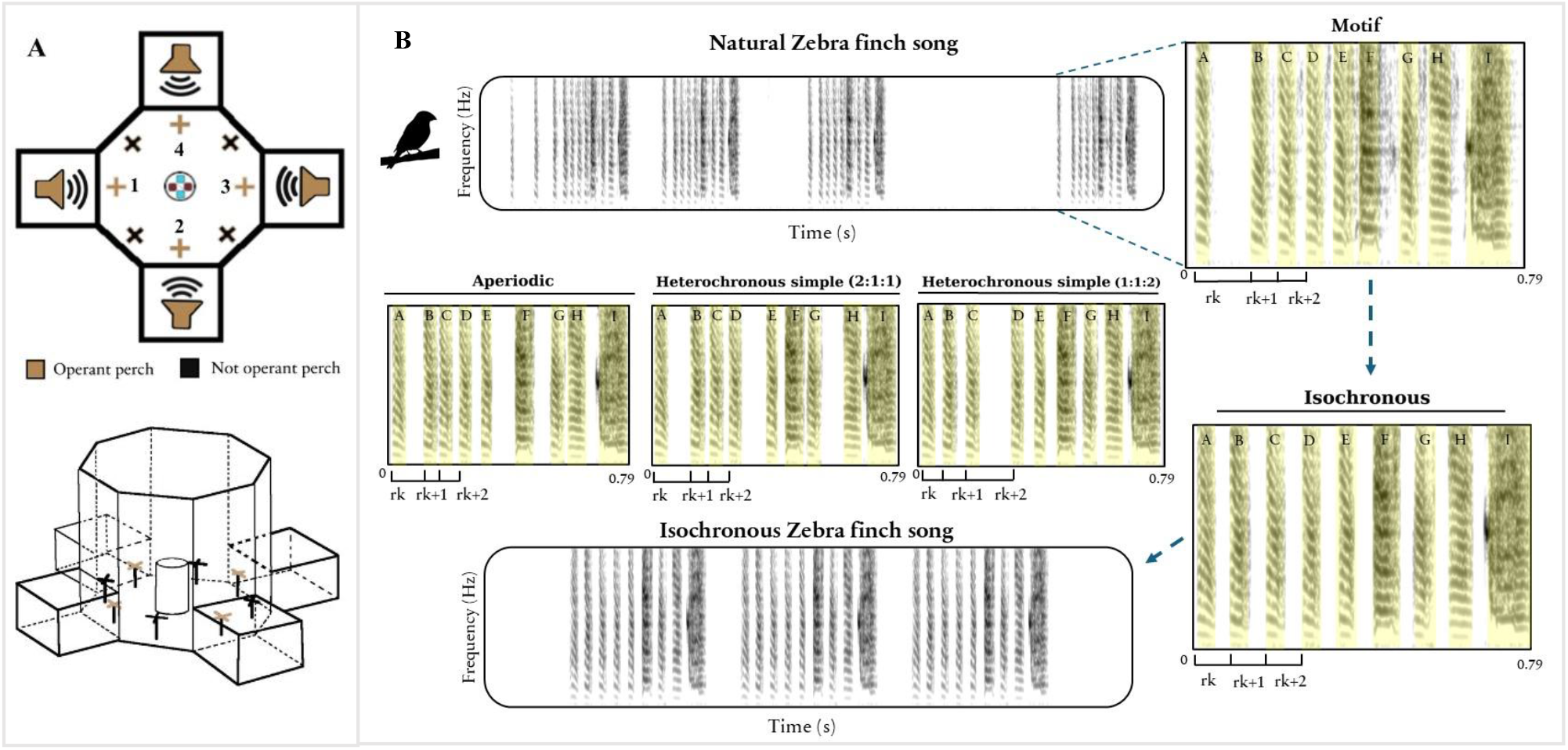
Operant preference setup and procedure used to construct the stimuli: **A)** Setup used to test preference for rhythmic patterns. Each perch, numbered from 1 to 4, elicited the sound from the nearby speaker. Food and water were available *ad libitum* at the centre of the arena. **B**) Procedure used to construct the stimuli. An edited motif was created from the original zebra finch song motif. The edit consisted of moving the syllables to a grid fitting the required categorical rhythm, maintaining the order and the spectral features of the syllables (A-I). From the edited motif (i.e. isochronous), the song with the categorical rhythm was formed by concatenating three motifs separated by 200 ms of silence. The four different edits were fitting the isochronous rhythm, the aperiodic rhythm and two versions of the heterochronous simple rhythm (2:1:1 & 1:1:2).

### Stimuli

Thirteen zebra finch song motifs were selected to create the experimental stimuli. The motifs belonged to 13 different zebra finches’ songs recorded at Leiden University from 2017 to 2021 and were edited to fit four different categorical rhythmic patterns. The motifs were considered suitable for editing if they contained more than three syllables and an odd number of syllables per motif. The original motifs were cut from the original song, and their syllables were annotated using the annotate-text grid function in Praat (version 6.4.06) [55], measuring the motif’s total duration, the temporal intervals between the syllables, and extracting its rhythmic patterns. From the original motif, four edited versions were generated: the isochronous version (1:1), the aperiodic version (random), and two heterochronous simple versions with the longer interval positioned either at the beginning (2:1:1) or at the end of the pattern (1:1:2). The syllables retained their spectral features, sequential order, and total duration as in the original motif. The edited version was modified only in the silence gaps between the syllables to match the categorical rhythmic pattern with a fixed protocol. We created a grid for each new motif, with the number of gridlines matching the number of syllables in the original motif. The gridlines were positioned to produce the intended rhythmic pattern, while preserving the total duration of the original motif. For example, for a pattern 1:1:2 and a motif containing 9 syllables, the sequence would have 9 gridlines matching the syllable onsets. The duration of the repeated section (1:1:2) was calculated by dividing the motif duration by the ratio of the number of syllables (9) to the number of elements in the repeated section (3). The duration of each grid interval was consequently derived from the duration of the repeated section (1:1:2). For the aperiodic pattern, which does not have any repetition, a random seed was generated in Excel, fitting those values to the number of required gridlines (Figure 1B). Humans recognise isochronous patterns more readily than zebra finches [39,56-58] but cannot perceive deviations from an isochronous pattern if they are less than 4% of the interval [59,60]. Therefore, from the perfect inter-onset interval, we calculated a 4% range to obtain an edited song that constitutes a natural rhythmic alternative to the zebra finch song and not an artificial version of it with a perfect rhythmic pattern, which is unlikely to be found in nature [18-22,61]. In some generated motifs, syllable overlap arose as a byproduct of differences in syllable duration. The 4% range helped to mitigate this effect by adjusting syllables to avoid overlap when possible while maintaining the overall rhythmic pattern. The overlap between syllables was present in 40% of motifs used, ranging from 15% of the stimuli in the aperiodic rhythm to 69% of the heterochronous simple stimuli (2:1:1) (Electronic supplementary material, Figure S6A-B, Table S6). Where the overlap occurred, its duration ranged from 1 to 155 milliseconds (median = 20.5 milliseconds). The final stimulus songs were made in PRAAT and composed by combining the edited motif three times, separated by 200 milliseconds of silence. To be certain that each motif would represent the rhythmic pattern desired, we checked the rhythmicity using the method of Roeske and colleagues [50] and visualised the patterns using the ternary plots [4,62-63] (Figure 2).

**Figure 2.**
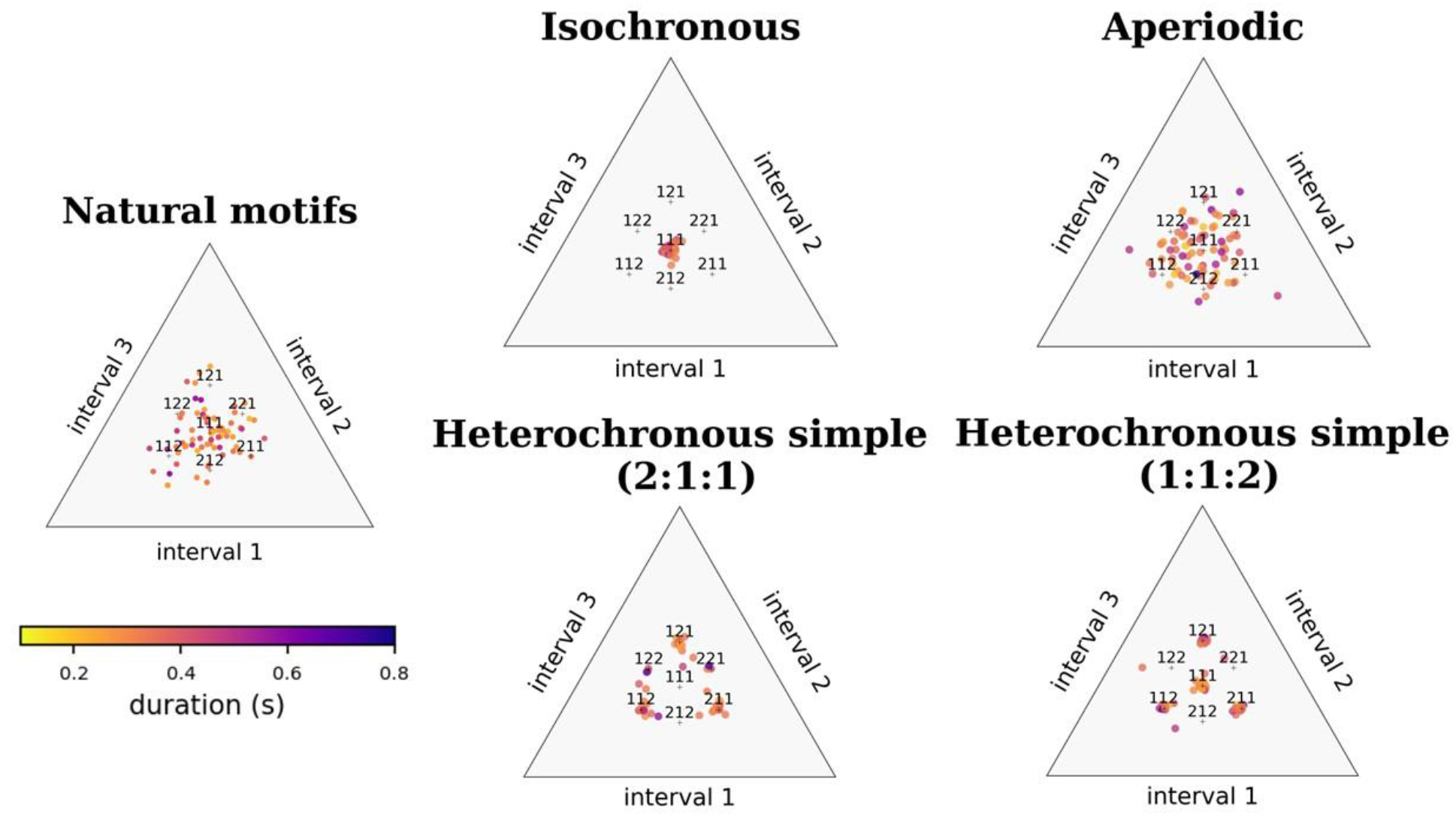
Categorical rhythmic patterns of the stimuli used: The dots represent the intervals (IOIs) within each stimulus, all pooled together. On the left, for comparison, the IOIs of the natural motifs of zebra finches were represented, which were not used as experimental stimuli.

### Procedure

The first experiment consisted of a 2-choice test with the isochronous and the aperiodic songs; the second experiment consisted of a 4-choice test with songs of all four rhythmic patterns (isochronous, aperiodic, heterochronous simple (2:1:1), heterochronous simple (1:1:2). Two testing birds and a guide bird of the same sex were moved from the home aviary into the operant setup for a day. In this phase, no sound was triggered when landing on the perches, but the birds habituated and explored the arena. On the following day, the guide bird and testing bird 2 were removed from the setup, allowing the other bird to be tested individually. This procedure was repeated for each bird, such that they were tested twice in each experiment. An experiment lasted 8 hours (9:00 – 17:00) and consisted of two 4-hour blocks (9:00-13:00; 13:00-17:00), with the sounds being automatically switched between the blocks and associated with the opposite perch (1 <-> 3; 2 <-> 4). The association between the perch number and the rhythmic pattern was pseudo-randomised at the start of the experiment, ensuring 4 blocks with different perch-sound associations for each bird. This counterbalancing procedure was implemented to minimise positional biases in the rhythmic preference data. At the end of the experiment, the guide bird and testing bird 2 were again moved into the setup. The next morning, the guide bird and testing bird 1 were removed from the setup, starting the experiment with testing bird 2. Each bird repeated the same choice test 1–3 days after the first test day. During the period between experiments (from 17:00 to 9:00), no sound could be triggered by landing on the perches. For the whole duration of the experiment, a background sound of their home aviary was played at around 50 dB from a JBL Clip+ speaker. In the 2-choice test, perches 1 and 3 were associated with the isochronous and the aperiodic version of the original motif, while perches 2 and 4 did not trigger any sound. In the 4-choice test, perches 1 and 3 were associated with the heterochronous simple rhythmic patterns (2:1:1 & 1:1:2), while perches 2 and 4 were associated with the isochronous and the aperiodic rhythmic patterns.

### Statistical analysis

Of the thirty-one birds tested in the operant preference setup, twenty-seven (17 females and 10 males) actively participated in at least one of the two preference experiments. From each experimental block, we recorded the number of times a bird triggered the sound (*original count*). We then extracted the number of times a bird triggered each operant perch after it had heard all sounds presented in that block (*count*). To ensure that birds had sufficient experience with all available response options, we included only blocks in which each operant perch had been selected at least once. We additionally applied an inclusion criterion equal to the number of operant perches * 2, resulting in a minimum of 4 responses for the 2-choice test and 8 responses for the 4-choice test. Based on these criteria, from the 27 birds tested in at least one of the experiments, data from 20 birds (10 females and 10 males) were retained for the 2-choice test, and data from 19 birds (10 females and 9 males) were retained for the 4-choice test.

Because birds made multiple choices in the same experimental block, responses were not treated as independent observations. We therefore analysed the distribution of responses among the available choices using multinomial choice models implemented with the mclogit package in R (*mclogit* function of mclogit package) [64,65]. Each combination of bird, testing date, and experimental block defined a multinomial choice set (*set*). Thus, the counts associated with the different rhythmic patterns and perch positions within a given set represented alternative outcomes of the same choice period.

For each experiment, we fitted multinomial models with rhythmic pattern and perch position as fixed effects. Syllable overlap and sex were additionally included in alternative models to assess their contribution to response choice. Bird identity was included as a random effect, allowing responses to rhythmic pattern to vary among individuals. Song motif was not included as a random effect because each bird was presented with the same motif throughout its experiment; consequently, motif was held constant within bird while the rhythmic pattern of that motif was experimentally manipulated. The different candidate models were compared using Akaike’s Information Criterion (AIC), and the model with the lowest AIC was selected as the best-supported model. Pairwise comparisons among rhythmic patterns were obtained using estimated marginal means with the emmeans package in R (*emmeans* package [66]). We used the same multinomial modelling approach to assess sex-level rhythmic pattern preferences, including an interaction between sex and rhythmic pattern. We also assessed positional preferences by fitting a multinomial model with perch position as the fixed effect.

To assess whether preference for a specific rhythmic pattern differed between the 2-choice test and 4-choice test, we additionally fitted a combined multinomial model including experiment type, rhythmic pattern, and their interaction, while controlling for perch position and including bird identity as a random effect. The interaction between test type and rhythmic pattern was used to assess whether responses to the rhythmic patterns differed between the 2-choice and 4-choice experiments.

Finally, to examine individual-level deviations from chance, we calculated for each bird the proportion of responses directed towards the isochronous stimulus. We tested whether this proportion differed from the expected probability using a binomial test [67], with a null probability of 0.50 for the 2-choice test and 0.25 for the 4-choice test.

## Results

### Rhythmic preferences

#### 2-choice test

In the 2-choice test, we found no significant overall preference for either isochronous or aperiodic rhythmic patterns (multinomial logistic regression with random slopes for rhythmic pattern by bird identity; Rhythmic pattern isochronous: β = −0.026, SE = 0.327, z = −0.08, p = 0.936). The estimation of marginal means indicated equivalent choice probabilities for both rhythmic patterns (Aperiodic: 0.468 ± 0.362 SE; Isochronous: 0.441 ± 0.434 SE; contrast: β = 0.027, SE = 0.336, t = 0.081, p = 0.936) (Electronic supplementary material, Figure S1, Table S1).

The overlap between syllables, which emerged as a byproduct of the procedure, did not significantly influence the choice between the different rhythmic patterns (Overlap: β = 0.009, SE = 0.722, z = 0.013, p = 0.990). There were no significant differences in rhythm preference between male and female zebra finches (*Female*, contrast Aperiodic – Isochronous: β = 0.065, SE = 0.471, t = 0.138, p = 0.891; *Male*, contrast Aperiodic – Isochronous: β = −0.010, SE = 0.456, t = - 0.021, p = 0.983) (Electronic supplementary material, Table S2). Overall, these results indicate that zebra finches do not show either a general preference or a sex-specific preference for isochronous or aperiodic rhythmic patterns (Figure 3).

**Figure 3.**
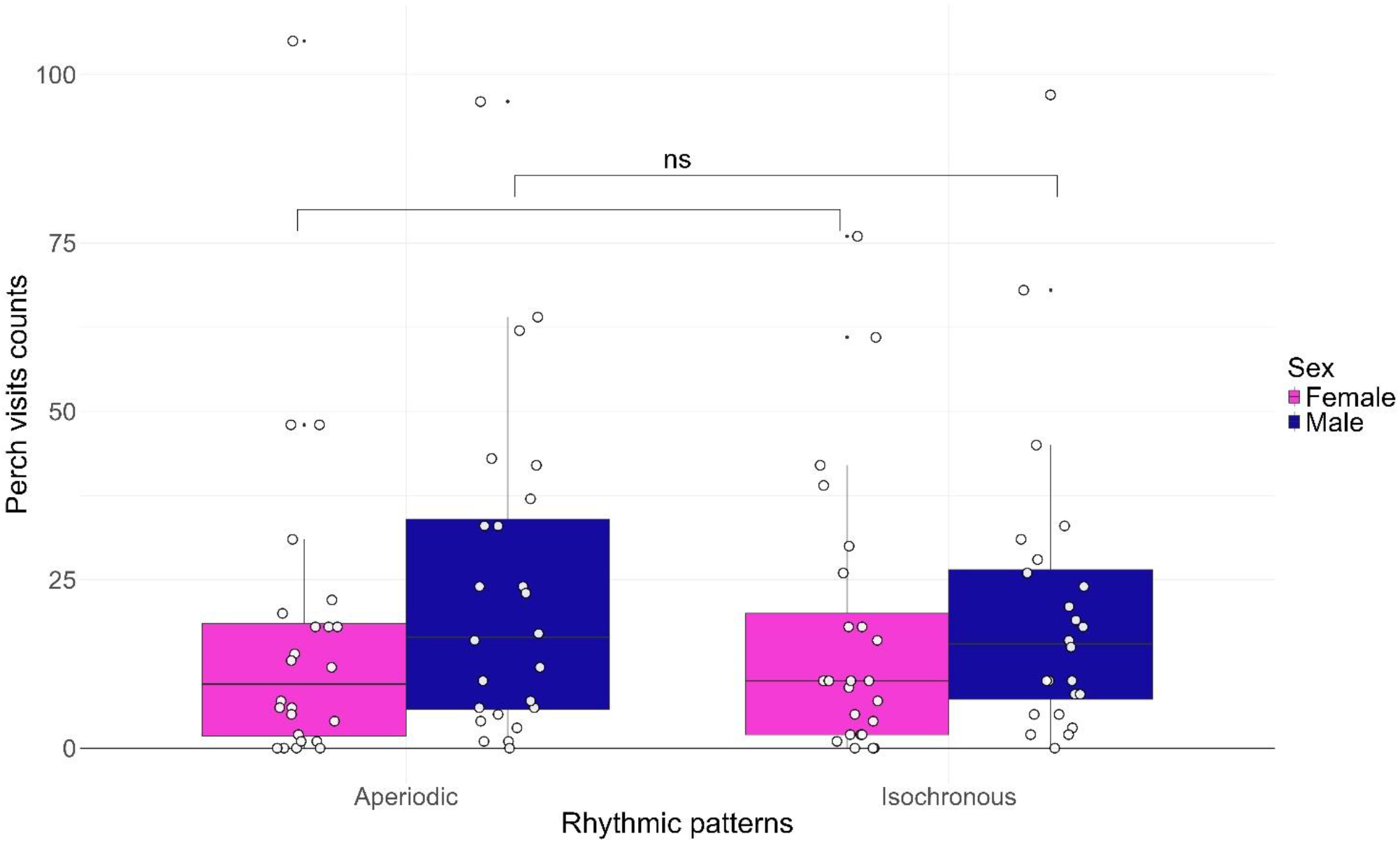
Sex preference for categorical rhythmic patterns in the 2-choice test: Data points represent the number of landings per block on perches triggering Aperiodic or Isochronous stimuli. Counts were extracted after birds had heard all stimuli presented within the experimental block and met the inclusion criterion of at least three responses. Responses are shown separately by sex. An experimental block was a 4-hour period during which each rhythm was associated with a unique perch position. Rhythmic patterns were counterbalanced across experimental blocks.

#### 4-choice test

In the 4-choice test, we found that the heterochronous simple rhythmic pattern ending with a long interval (1:1:2) was visited less (β = −0.645, SE = 0.234, z = −2.751, p = 0.006) than the aperiodic one. The isochronous rhythmic patterns did not differ from aperiodic patterns (β = −0.025, SE = 0.262, z = −0.097, p = 0.923), confirming the findings in the 2-choice test (Electronic supplementary material, Table S3, Figure S3). The overlap between syllables, contrary to the results of the 2-choice test, played a significant role in the selection of the song (Overlap: β = −0.677, SE = 0.265, z = −2.552, p = 0.011). However, it is significant only in the 1:1:2 heterochronous simple rhythmic pattern when looking at the interaction between rhythmic pattern and overlap (Electronic supplementary material, Table S7, Figure S6).

Females did not show a preference for any of the rhythmic patterns (Aperiodic – Heterochronous simple (1:1:2): β = 0.817, SE = 0.336, t = 2.429, p = 0.078; Aperiodic – Heterochronous simple (2:1:1): β = 0.386, SE = 0.323, t = 1.195, p = 0.632, Aperiodic - Isochronous: β = -0.128, SE = 0.342, t = -0.373, p = 0.982). In contrast, males showed a preference for the aperiodic rhythmic pattern compared to both heterochronous simple rhythmic patterns (Aperiodic – Heterochronous simple (1:1:2): β = 1.042, SE = 0.353, t = 2.954, p = 0.020; Aperiodic –Heterochronous simple (2:1:1): β = 1.184, SE = 0.345, t = 3.432, p = 0.005) but not compared to the isochronous rhythmic pattern (Aperiodic - Isochronous: β = 0.342, SE = 0.363, t = 0.944, p = 0.781) (Electronic supplementary material, Table S4) (Figure 4). These results point to a lack of general preference for a rhythmic pattern in zebra finches, with males selecting the aperiodic rhythmic pattern more than the heterochronous simple ones. At the same time, the males do not present any significant preference when comparing the aperiodic with the isochronous rhythmic pattern, similar to the results in the 2-choice test.

**Figure 4.**
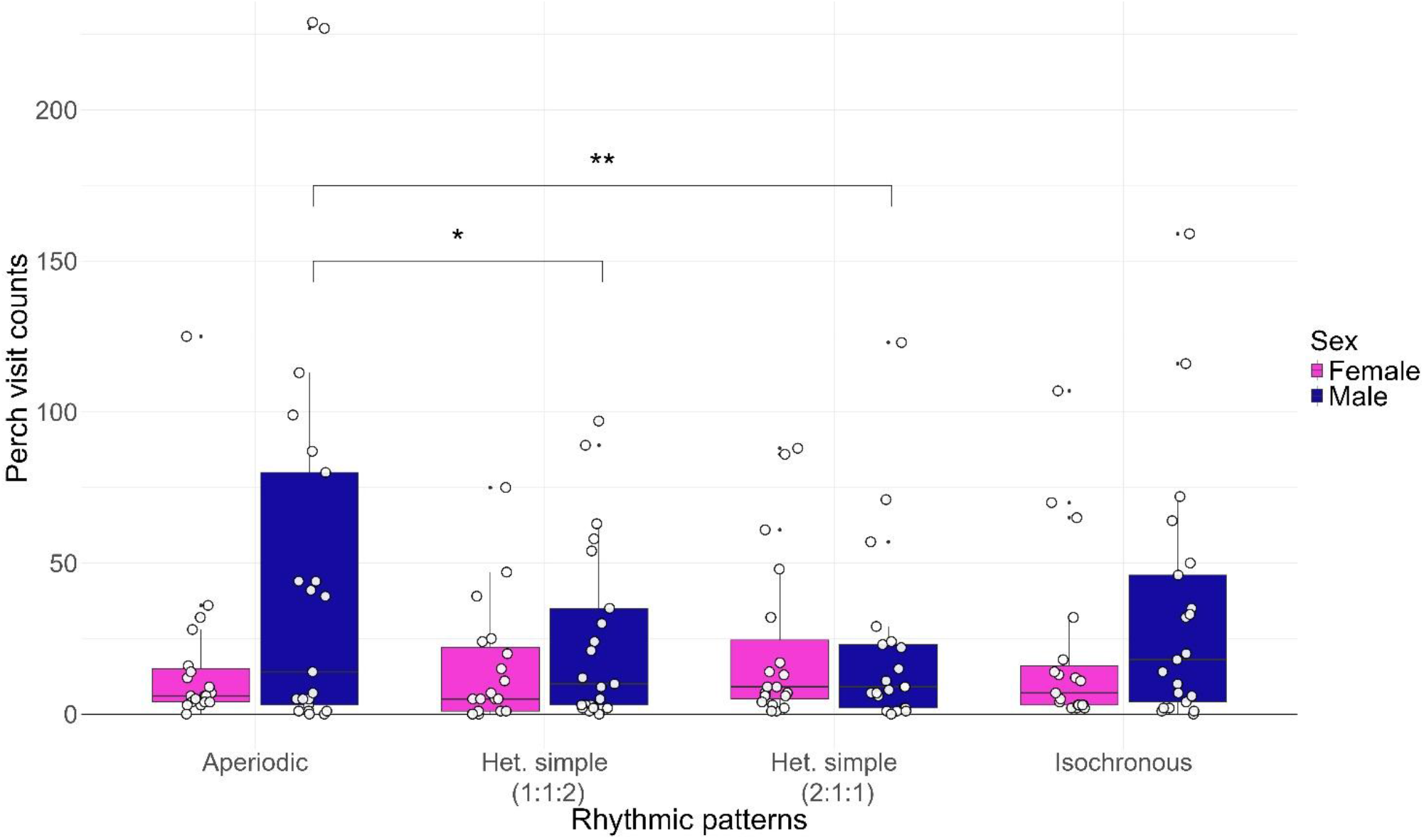
Sex preference for categorical rhythmic patterns in the 4-choice test: Data points represent the number of landings per block on perches triggering Aperiodic, Heterochronous simple (1:1:2), Heterochronous simple (2:1:1) or Isochronous stimuli. Counts were extracted after birds had heard all stimuli presented within the experimental block and met the inclusion criterion of at least five responses. Responses are shown separately by sex. An experimental block was a 4-hour period during which each rhythm was associated with a unique perch position. Rhythmic patterns were counterbalanced across experimental blocks.

#### Positional preferences

Across both tests, zebra finches showed significant preferences for particular perches, independent of the experimental condition. In the 2-choice test, perch 3 received more visits than perch 1 (β = 0.928, SE = 0.061, z = 15.19, p < 0.001). In the 4-choice test, the same preference was observed, with perch 3 receiving more visits than perch 1 (β = 0.138, SE = 0.059, z = 2.34, p = 0.019), and with perch 2 generally avoided compared to perch 1 (β = −1.548, SE = 0.059, z = −26.41, p < 0.001). The avoidance of perch 2 and perch 1 was consistent with their position in the setup, being the closest perches to the door of the room. A multinomial model with only perch position as the response variable showed a lower AIC (AIC = 614.64) than the model with rhythmic stimuli (AIC = 1145.89), indicating perch number as one of the main factors explaining the variance in the data and suggesting a positional preference in the arena (Electronic supplementary material, Figure S5, Table S5).

### Individual differences in rhythmic preference

To assess whether the population preference for a rhythmic pattern remained stable across the two experiments, we calculated the preference ratio for the isochronous rhythmic pattern. Among the 20 birds tested, 13 birds participated in both the 2-choice and 4-choice tests. The population preference for the isochronous rhythmic pattern (2-choice test mean = 0.465 ± 0.239 SD; 4-choice test mean = 0.200 ± 0.198 SD), dropped in the 4-choice test compared to the 2-choice test (paired t-test: t_12_ = 3.02, p = 0.011, mean difference = 0.265 ± 0.079 SD), suggesting that there was an influence of the number of different rhythmic patterns available, rather than a change in preference between the isochronous and aperiodic patterns (Test × Isochronous: β = 0.112, SE = 0.091, z = 1.24, p = 0.216).

Although at the population level we did not find a general preference for rhythmic patterns, individual birds showed preferences that deviated significantly from chance: 9 of 20 birds in the 2-choice test and 12 of 19 birds in the 4-choice test selected rhythmic patterns over the others consistently (binomial test, p < 0.05 per bird). This proportion far exceeded the ∼5% (α = 0.05) expected if no birds truly differed from chance (binomial test on the number of significant birds: 2-choice test, p = 1.98 × 10^−7^; 4-choice test, p = 8.84 × 10^−12^), indicating variation on rhythm selection among individuals, rather than no preference for any rhythmic pattern at the individual level (Electronic supplementary material, Figure S2 and S4).

## Discussion

In this study, we investigated whether zebra finches showed a preference for categorical rhythmic patterns. Zebra finches did not show convergence on a shared preference for any categorical rhythmic pattern, regardless of the number of alternatives available (2-choice and 4-choice tests). However, in the 4-choice test, birds avoided the heterochronous simple rhythmic pattern with a long interval at the end of the sequence (1:1:2) and the songs containing overlapping syllables. Because overlap significantly reduced responses and occurred more frequently in the heterochronous rhythmic stimuli, it may have contributed to the observed avoidance of heterochronous simple rhythmic patterns (1:1:2). Nevertheless, syllable overlap was most common in the heterochronous simple stimuli with a longer interval at the start of the sequence (2:1:1) (Electronic supplementary material, Figure S6A, Table S6-S7), which did not exhibit the same pattern of avoidance, suggesting that syllable overlap alone is unlikely to fully explain the effect. No differences in preferences were observed between male and female zebra finches in the 2-choice test. In contrast, in the 4-choice test, only males avoided heterochronous simple rhythmic patterns, whereas females showed a weaker, non-significant trend in the same direction. This pattern indicates that males primarily drove the overall avoidance of heterochronous rhythmic patterns. In both tests, responses varied substantially between individuals, without any rhythmic category emerging as consistently preferred at the population level. Finally, birds showed a consistent spatial bias, preferring perches located furthest from the room entrance.

Our findings suggest that zebra finches do not have a preference for songs with any of the categorical rhythmic patterns tested. Although only males sing, both sexes show preferences for conspecific songs [68-70]. The songs preferred by females in auditory choice tasks predict mate choice preferences in a social setting [71]. Males also tend to use particularly preferred songs to attract conspecifics [42,70]. Numerous studies have examined which features of the male song are the most important for song preference in both females and males, with a particular focus on the spectral information [69,72-73]. This study suggests that categorical rhythmic patterns alone do not play an important role in song selection and therefore may not substantially influence mate choice preferences. However, it does not imply that zebra finches do not pay attention to the temporal information of the song, as they showed an avoidance of the songs with overlapping syllables, suggesting a response toward songs that cannot be produced naturally. During the 4-choice test, zebra finches showed a tendency to visit heterochronous simple rhythmic patterns (1:1:2) less than aperiodic rhythmic patterns. However, the same tendency was not observed with the heterochronous simple rhythmic pattern with a long interval at the start (2:1:1), which may suggest that zebra finches do not perceive those discrete patterns as belonging in the same category, but may avoid specific temporal intervals or sequences.

Similarly, a lack of preference at the population level was found in budgerigars for sound strings with a regular or irregular beat [16]. Even though the aims of the two studies were different, these findings suggest the absence of a population bias toward the aperiodic and periodic embedded rhythmic patterns in a string of sounds. Recent work by Haiduk and colleagues [74] further inspected whether budgerigars prefer certain rhythmic patterns, using an operant preference setup. In this study, they investigated whether rhythmic preference in budgerigars is correlated with regularity, defined as sonic events fitting with categorical rhythmic patterns, or with repetition, defined as recurrence of patterns over a sequence. Whilst the study did not aim to investigate the preference among categorical rhythmic patterns, isochronous and heterochronous rhythmic patterns were included in the stimuli used, making a direct comparison between the two studies possible. In accordance with the earlier study, testing regular and irregular beat preference in budgerigars, and our findings on zebra finches, no population-level preference for a rhythmic pattern was found. However, in agreement with previous findings on budgerigars and in contrast to our findings on zebra finches, sex-specific preferences were reported, with female budgerigars choosing sounds with either pattern repetitions or regularity over silence. The authors indicate that both regularity and repetition are important aspects in rhythm perception in budgerigars, suggesting that the sex-specific preference in budgerigars could be driven by complex rhythmic vocal displays in the species, such as language in humans and warble song in budgerigars, which share similar rhythmic features [75].

In this framework, zebra finch song has been reported as isochronous in two different studies [49,50]. However, even though the study of Norton & Scharff reports a repetition of temporal intervals in the song that they call metronomic, those temporal intervals were not quantified, and therefore, a direct link with isochrony is missing. Roeske and colleagues inspected the zebra finch song rhythm and the presence of categorical rhythms, finding a non-significant normal distribution around isochrony. These findings suggest isochrony more as the mean of different rhythms [76] rather than the predominant rhythmic pattern in the song. A similar distribution is evident in our visualisation of the original motif of the zebra finch (Figure 2), where the inter-onset intervals range around isochrony, without fitting it. Recent work on the zebra finch’s song seems to corroborate this evidence, assessing the song of zebra finches as not generally isochronous, but with an isochronous section nested in it [77,78]. Together, these observations suggest that zebra finches may be sensitive to temporal structures that are characteristic of their own vocalisations rather than to the categorical rhythmic patterns commonly described and found in human music.

Birds showed a strong positional preference across both experiments, with perches located furthest from the room entrance generally receiving more visits. Although the operant setup was positioned at the centre of the room and perch-song associations were counterbalanced across blocks, the position of the door between perches 1 and 2 likely influenced spatial use within the arena. Importantly, because rhythmic stimuli were systematically switched between opposite perch locations across blocks, this bias is unlikely to account for the absence of general rhythmic preference.

Together with evidence from budgerigars, our results suggest that preferences for categorical rhythmic patterns are not a universal feature amongst vocal-learning species. In humans, however, categorical rhythmic patterns are widespread across cultures, emerge spontaneously in transmission experiments, and are often associated with perceptual preference [4,10,16]. This contrast suggests that preferences for categorical rhythmic patterns may be human-specific and may be connected with species-specific vocal displays.

In conclusion, zebra finches responded differently to the different temporal organisation of the songs but did not converge on a shared preference for any categorical rhythmic pattern. Nevertheless, they avoided songs containing overlapping syllables and, in some contexts, heterochronous simple rhythmic patterns. These findings suggest that temporal organisation may influence song preferences in zebra finches, but that the categorical rhythmic patterns characteristic of human music do not appear to be a primary driver of song preference in this species.

