## Supplementary Material for "Categorical rhythmic patterns do not predict song preference in zebra finches"

Supplementary Materials for: **Categorical rhythmic patterns do not predict song preference in zebra finches**

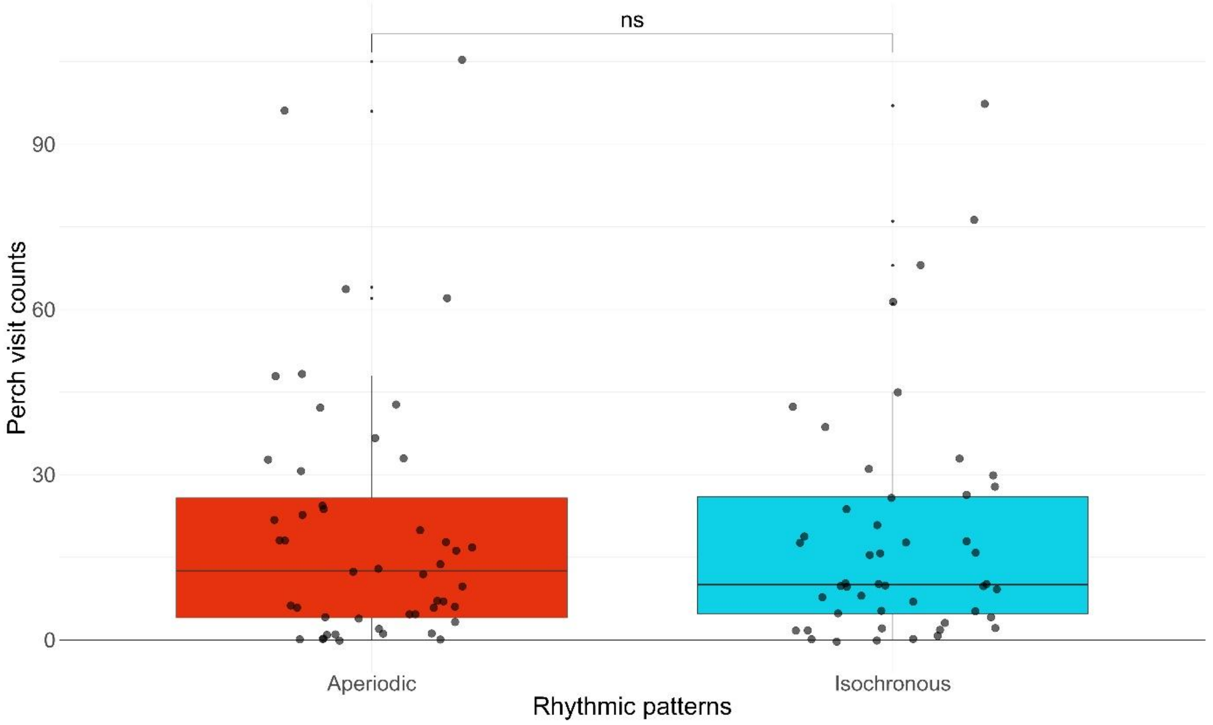


**Figure S1| General preference for categorical rhythmic patterns in the 2-choice test:** Data points represent the number of landings per block on perches triggering Aperiodic or Isochronous stimuli. Counts were extracted after birds had heard all stimuli presented within the experimental block and met the inclusion criterion of at least three responses. An experimental block was a 4-hour period during which each rhythm was associated with a unique perch position. Rhythmic patterns were counterbalanced across experimental blocks.

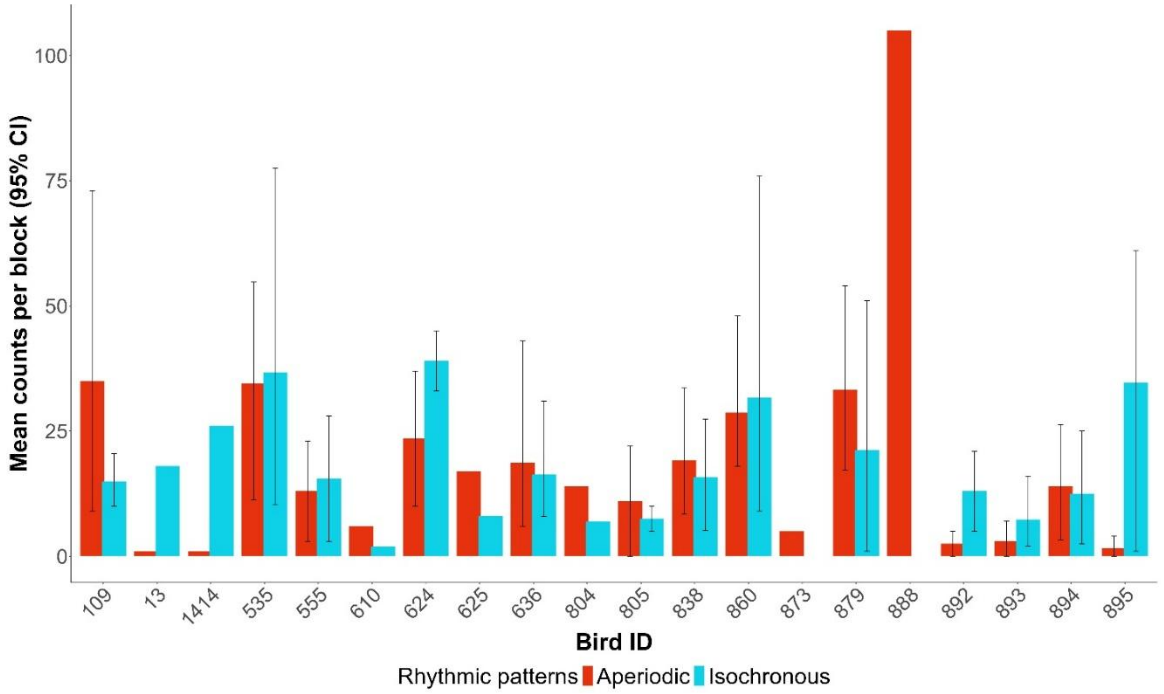


**Figure S2| Individual preference for categorical rhythmic patterns in the 2-choice test:** The graph represents the individual preference per bird in the 2-choice test. Each bar plot shows the mean counts per block (1-4 blocks), with a block being a 4-hour session with a fixed association between rhythmic patterns and perch position. Rhythmic patterns – perch position were counterbalanced over blocks.

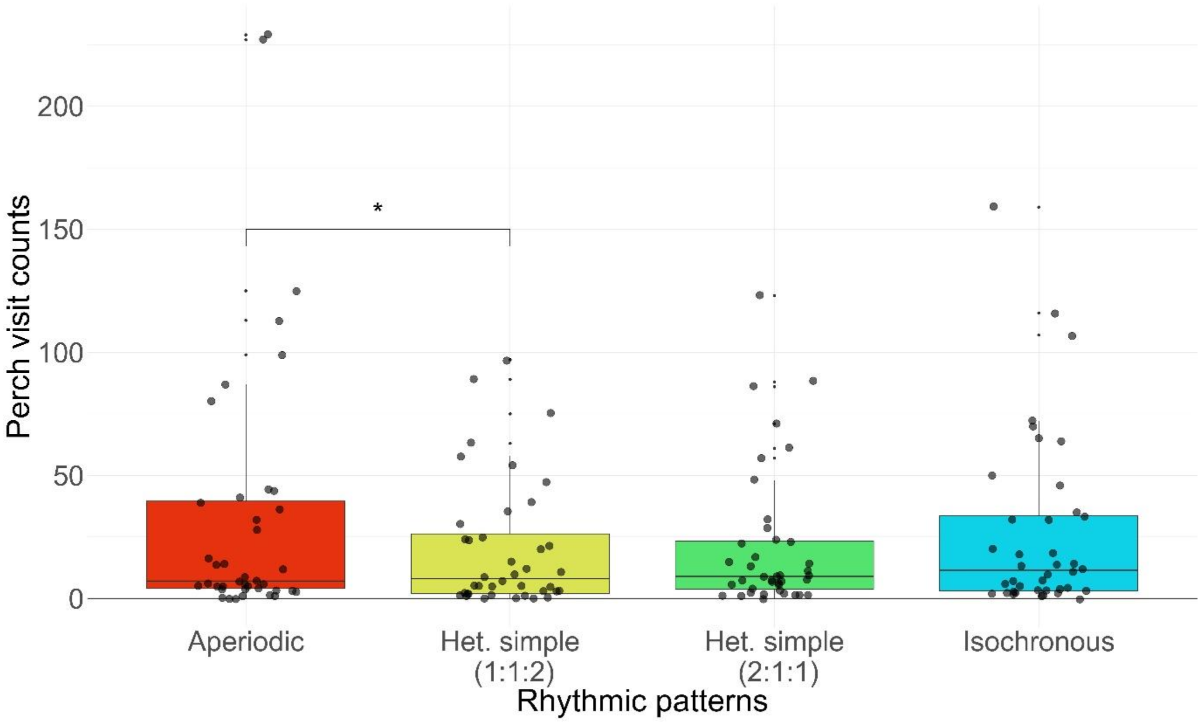


**Figure S3| General preference for categorical rhythmic patterns in the 4-choice test:** Data points represent the number of landings per block on perches triggering Aperiodic, Heterochronous simple (1:1:2), Heterochronous simple (2:1:1) or Isochronous stimuli. Counts were extracted after birds had heard all stimuli presented within the experimental block and met the inclusion criterion of at least five responses. An experimental block was a 4-hour period during which each rhythm was associated with a unique perch position. Rhythmic patterns were counterbalanced across experimental blocks.

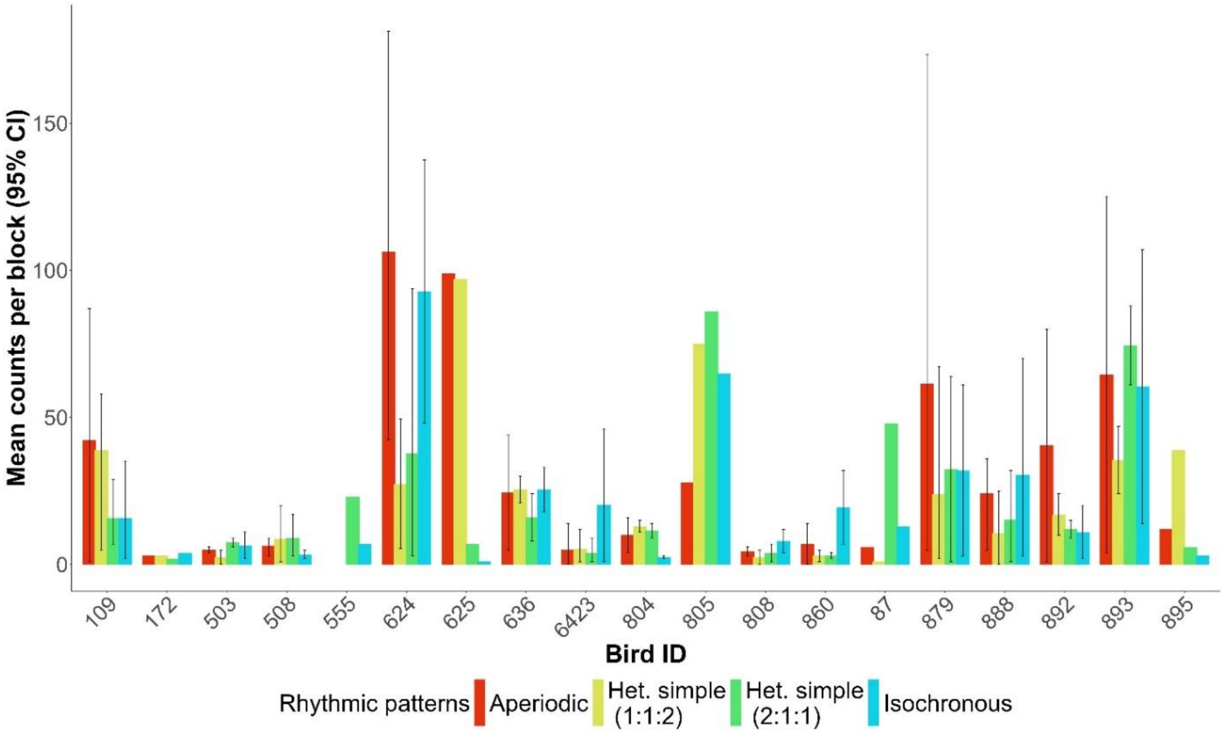


**Figure S4| Individual preference for categorical rhythmic patterns in the 4-choice test:** The graph represents the individual preference per bird in the 4-choice test. Each bar plot shows the mean counts per block (1-4 blocks), with a block being a 4-hour session with a fixed association between rhythmic patterns and perch position. Rhythmic patterns – perch position were counterbalanced over blocks.

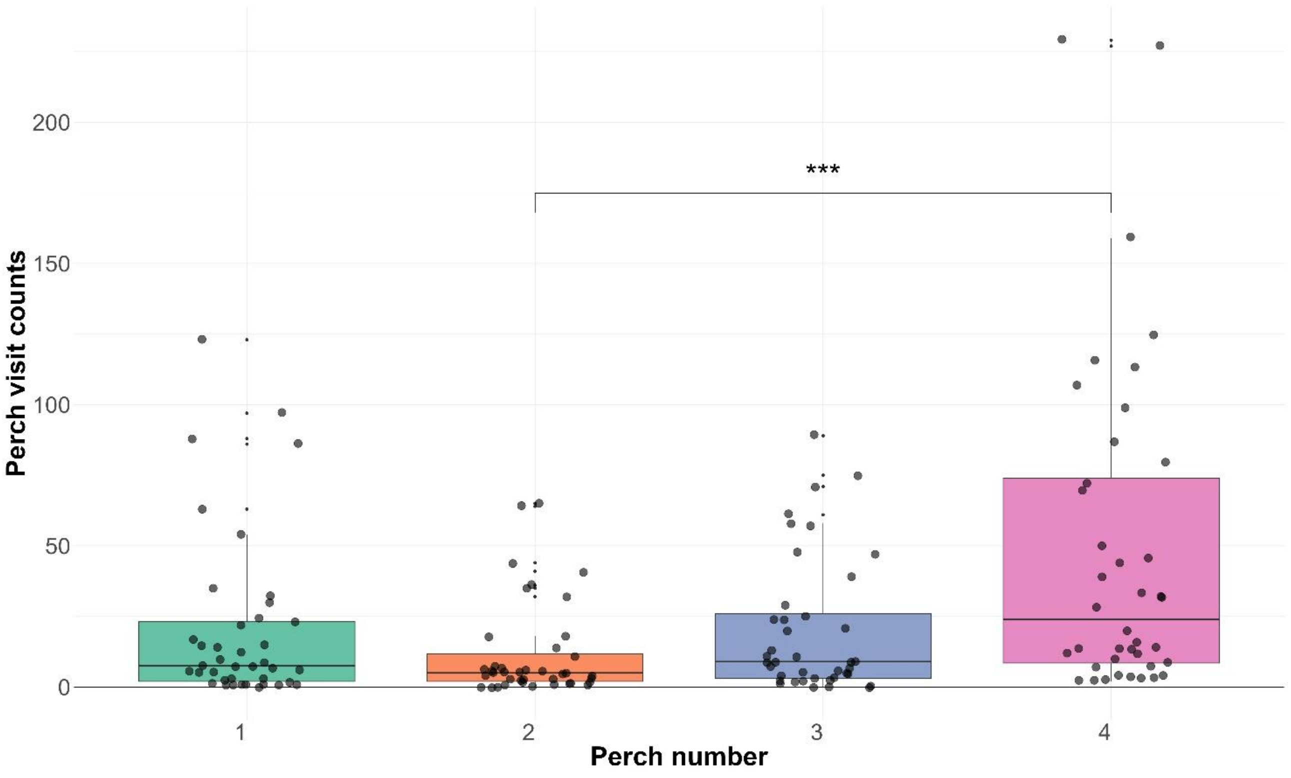


**Figure S5| General preference for perch position in the 4-choice test:** Data points represent the number of landings per block on perch positions (1,2,3,4). Counts were extracted after birds had heard all stimuli presented within the experimental block and met the inclusion criterion of at least eight responses for the 4-choice test. An experimental block was a 4-hour period during which each rhythm was associated with a unique perch position. Rhythmic patterns were counterbalanced across experimental blocks. Perches 1-2 were the ones more close to the door to enter the room and 3-4 were the more distant (See Figure 1A).

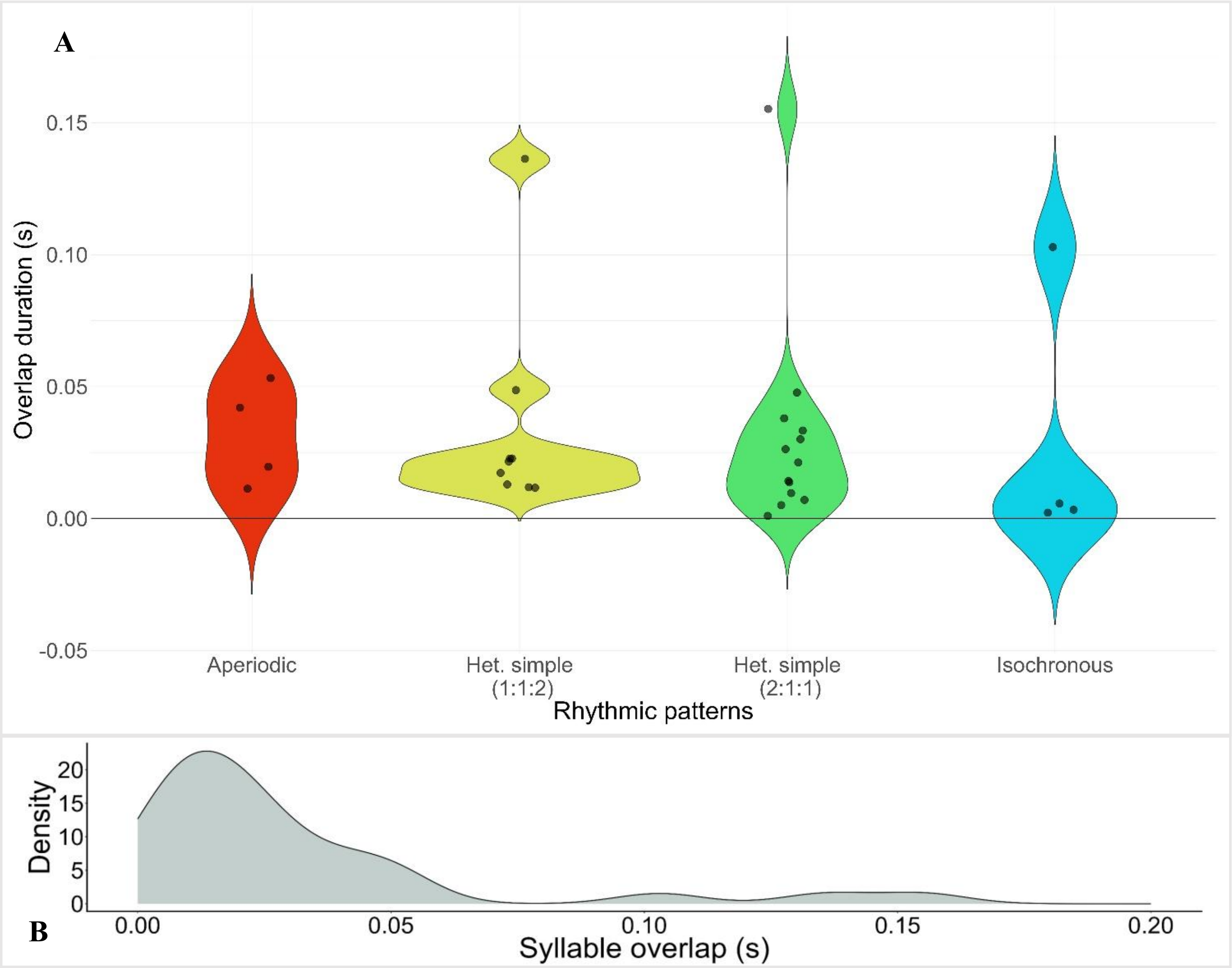


**Figure S6| Syllable overlap duration and distribution across the four categorical rhythms: A)** Presence of the syllable overlap over the four rhythmic patterns created. Syllable overlap was present in 40% of motif used, ranging from 15% of the aperiodic rhythmic pattern to the 69% of the heterochronous simple (2:1:1). **B)** Distribution of the syllable overlap across the four rhythmic patterns. The duration of the overlap ranged from 0.001 s to 0.155 s, with a median of 0.0205 seconds.

**Table S1| Results of the model for rhythmic patterns in the 2-choice test:** Outcome of the model used to compare aperiodic and isochronous selections across birds in the 2-choice test.

| contrast | estimate | SE | df | t.ratio | p.value |
|---|---|---|---|---|---|
| APE - ISO | 0.0272 | 0.336 | 47 | 0.081 | 0.9357 |

**Table S2| Results of the model with the interaction of rhythmic patterns and sex in the 2-choice test:** Outcome of the model used to compare aperiodic and isochronous selections for each sex in the 2-choice test.

| *Sex = Female:* | | | | | |
|---|---|---|---|---|---|
| **contrast** | **estimate** | **SE** | **df** | **t.ratio** | **p.value** |
| APE - ISO | 0.0649 | 0.471 | 47 | 0.138 | 0.8909 |
| *Sex = Male:* | | | | | |
| **contrast** | **estimate** | **SE** | **df** | **t.ratio** | **p.value** |
| APE - ISO | -0.0098 | 0.456 | 47 | -0.021 | 0.9829 |

**Table S3| Results of the model for rhythmic patterns in the 4-choice test:** Outcome of the model used to compare aperiodic, heterochronous simple (1:1:2), heterochronous simple (2:1:1) and isochronous selection across birds in the 4-choice test.

| contrast | estimate | SE | df | t.ratio | p.value |
|---|---|---|---|---|---|
| APE – HETSIMP (1:1:2) | 0.6449 | 0.234 | 108 | 2.751 | 0.0346 |
| APE - HETSIMP (2:1:1) | 0.3259 | 0.311 | 108 | 1.046 | 0.7226 |
| APE - ISO | 0.0253 | 0.262 | 108 | 0.097 | 0.9997 |
| HETSIMP (1:1:2) – HETSIMP (2:1:1) | -0.3189 | 0.289 | 108 | -1.104 | 0.6879 |
| HETSIMP (1:1:2) - ISO | -0.6195 | 0.300 | 108 | -2.067 | 0.1706 |
| HETSIMP (2:1:1) - ISO | -0.3006 | 0.263 | 108 | -1.141 | 0.6652 |

**Table S4| Results of the model with the interaction of rhythmic patterns and sex in the 4-choice test:** Outcome of the model used to compare aperiodic, heterochronous simple (1:1:2), heterochronous simple (2:1:1) and isochronous selections for each sex in the 4-choice test.

| *Sex = Female:* | | | | | |
|---|---|---|---|---|---|
| **contrast** | **estimate** | **SE** | **df** | **t.ratio** | **p.value** |
| APE – HETSIMP (1:1:2) | 0.817 | 0.336 | 106 | 2.429 | 0.0778 |
| APE - HETSIMP (2:1:1) | 0.386 | 0.323 | 106 | 1.195 | 0.6316 |
| APE - ISO | -0.128 | 0.342 | 106 | -0.373 | 0.9822 |
| 1:1:2 – 2:1:1 | -0.431 | 0.440 | 106 | -0.979 | 0.7616 |
| HETSIMP (1:1:2) - ISO | -0.944 | 0.406 | 106 | -2.325 | 0.0988 |
| HETSIMP (2:1:1) - ISO | -0.513 | 0.326 | 106 | -1.575 | 0.3973 |

| *Sex = Male:* | | | | | |
|---|---|---|---|---|---|
| **contrast** | **estimate** | **SE** | **df** | **t.ratio** | **p.value** |
| APE – HETSIMP (1:1:2) | 1.042 | 0.353 | 106 | 2.954 | 0.0199 |
| APE - HETSIMP (2:1:1) | 1.184 | 0.345 | 106 | 3.432 | 0.0047 |
| APE - ISO | 0.342 | 0.363 | 106 | 0.944 | 0.7810 |
| 1:1:2 – 2:1:1 | 0.141 | 0.468 | 106 | 0.302 | 0.9904 |
| HETSIMP (1:1:2) - ISO | -0.700 | 0.429 | 106 | -1.633 | 0.3649 |
| HETSIMP (2:1:1) - ISO | -0.841 | 0.347 | 106 | -2.427 | 0.0781 |

**Table S5| Results of the model for perch position in the 2 and 4-choice tests:** Outcome of the model used to compare the selection for perch 1, 2, 3 and 4 across birds and across the two tests (2-choice test, 4-choice test).

| contrast | estimate | SE | df | t.ratio | p.value |
|---|---|---|---|---|---|
| Perch.N1 - Perch.N2 | 0.710 | 0.333 | 111 | 2.132 | 0.1493 |
| Perch.N1 - Perch.N3 | 0.014 | 0.411 | 111 | 0.034 | 1.0000 |
| Perch.N1 - Perch.N4 | -0.714 | 0.284 | 111 | -2.511 | 0.0637 |
| Perch.N2 - Perch.N3 | -0.696 | 0.314 | 111 | -2.212 | 0.1262 |
| Perch.N2 - Perch.N4 | -1.424 | 0.340 | 111 | -4.183 | 0.0003 |
| Perch.N3 - Perch.N4 | -0.728 | 0.342 | 111 | -2.127 | 0.1508 |

**Table S6| Syllable overlap degree across the four different rhythmic patterns:** Analysis of the syllable overlap over the four rhythmic patterns; n is the number of syllable overlaps present across all the stimuli used, with the mean, median, and sd per rhythmic pattern. Min and Max are the minimum and the maximum value for each rhythmic pattern.

| Rhythmic pattern | n | mean | median | sd | min | max |
|---|---|---|---|---|---|---|
| Aperiodic | 4 | 0.0315 | 0.031 | 0.0194 | 0.011 | 0.053 |
| Heterochronous simple (1:1:2) | 9 | 0.0341 | 0.022 | 0.0399 | 0.012 | 0.136 |
| Heterochronous simple (2:1:1) | 13 | 0.0309 | 0.021 | 0.0398 | 0.001 | 0.155 |
| Isochronous | 4 | 0.0285 | 0.0045 | 0.0497 | 0.002 | 0.103 |

**Table S7| Results of the model for syllable overlap across the four rhythmic patterns:** Outcome of the model used to compare whether there is a significant difference between motifs without syllable overlap and motif with syllable overlap across the four rhythmic patterns.

| *Aperiodic rhythmic pattern* | estimate | SE | df | t.ratio | p.value |
|---|---|---|---|---|---|
| No Overlap - Yes Overlap | 0.166 | 0.507 | 105 | 0.327 | 0.7445 |
| *Het. simple rhythmic pattern (1:1:2)* | | | | | |
| No Overlap - Yes Overlap | 1.069 | 0.382 | 105 | 2.801 | 0.0061 |

| | | | | | |
|---|---|---|---|---|---|
| *Het. simple rhythmic pattern (2:1:1)* | | | | | |
| No Overlap - Yes Overlap | 0.564 | 0.462 | 105 | 1.220 | 0.2252 |
| *Isochronous rhythmic pattern* | | | | | |
| No Overlap - Yes Overlap | 0.465 | 0.484 | 105 | 0.959 | 0.3398 |
